# Cold stress upregulates the expression of heat shock and *Frost* genes, but the evolution of cold stress resistance is apparently not mediated through either heat shock or *Frost* genes in the cold stress selected populations of *Drosophila melanogaster*

**DOI:** 10.1101/2022.03.07.483305

**Authors:** Karan Singh, Nagaraj Guru Prasad

## Abstract

The ability to thermal adaptation can alter insects’ reproductive traits and physiologic response. A number of studies in *Drosophila* documented the higher expression levels of *heat shock proteins* (*Hsps*) and *Frost* (*Fst*) genes during the recovery phase of cold shock treatment and suggested that higher expression levels of these genes and lipids can protect from cold stress. However, these genes’ expression and cold adaptation have not been well studied. Therefore, understanding the molecular and biochemical basis of cold adaptation, we examined the levels of lipid and expression patterns of *Hsp22, Hsp23, Hsp40, Hsp68, Hsp70, Hsp 83*, and *Fst* genes under the cold stress or benign condition in the populations of *Drosophila melanogaster* selected for increased resistance to cold shock.

We observed the significant up-regulation of *Hsp22, Hsp23, Hsp40, Hsp68*, and *Fst* genes in the FSB and FCB populations during the recovery phase post cold shock compared to the no shock condition. However, there was not a significant change in the transcript levels of these genes between FSB and FCB under both the cold stress or no shock condition. Additionally, we noticed higher total lipid levels in the females from cold shock treatment than those from no shock treatment. This finding suggests that cold selected populations have different mechanisms to sustain cold stress.

## INTRODUCTION

Insects are present in diverse habitats ranging from hot springs to cold deserts. Insects are ectothermic organisms and, therefore, cannot regulate internal temperature when temperature changes in the external environment. Therefore, insects have evolved a number of mechanisms to protect themselves when subjected to external environmental fluctuations. Thermal stress (hot or cold) is known to affect multiple traits, including fecundity, egg viability, mating ability, and pre- and post-copulatory traits, and it also reportedly affects the physiology of the insect (reviewed by [1,2,3,4,5]. Multiple studies suggested that thermal stress causes damage to various molecules like proteins and lipids and damage to cell membranes, protein transport machinery [6,7,8,9]. Insects are known to have evolved a range of mechanisms to mitigate and prevent the damage caused due to cold or heat stress. Change in lipid composition imparting more fluidity to cell membranes is one of the significant changes that is often employed [10].

Multiple proteins involved in repairing protein damage, including misfolding, aggregation, and denaturation are, known as heat shock proteins, play a crucial role. Heat shock proteins (Hsps) mitigate the effects of heat stress by protecting proteins and enzymes and facilitating their proper functioning under stressful conditions [11,12]. Also, an increase in the synthesis of several other metabolites like glycerol [13], glycogen, and cryoprotectants like trehalose proline. These metabolites help insects to survive under thermal stress [14]. Various insects like flesh fly [15], stick insects [16], field crickets [17], and corn borer [18], amongst others, have been used to study the mechanisms that are involved in coping with environmental stress. These insects employ a number of mechanisms that could be distinct yet overlapping. *D. melanogaster* has been a favorite model to address questions pertaining to underlying mechanisms of cold tolerance. Results from these studies suggest that *Drosophila* also employs mechanisms like lipid modification, increase in expression of heat shock proteins, synthesis of other metabolites like proline, glycogen, etc. to survive under thermal stress [13,7,19,20].

However, the expression pattern of *Hsps* and *Fst* genes has not been well studied in the evolved population against cold stress to understand the molecular basis of cold adaptation. To assess the molecular and biochemical mechanism of evolved cold stress resistance, we used cold shock selected populations of *Drosophila melanogaster* (*D. melanogaster*) that has evolved for a number of traits, including better survival, increased egg viability, more mating frequency, higher male mating ability, and ability to sire more progeny, we measured the total lipid content and transcripts levels of *Hsp22, Hsp23, Hsp40, Hsp68, Hsp70Aa* (*Hsp70*), *Hsp83*, and *Fst* genes under the cold stress or no shock condition.

## MATERIAL AND METHODS

Maintenance and derivation details of the experimental populations (FSB and FCB) are described previously [1,2,3,4,5] Experimental flies were generated from standardized flies, as explained previously in [1,2]. Briefly, flies from both FSB and FCB regimes were not subjected to any selection for one generation. Eggs were collected from these standardized flies at a controlled density of 70 eggs per vial. Eighteen vials were established for each of the FSB and FCB populations. Flies that emerged from these vials were used for the experiments. Gene expression was measured only in males, while both males and females were used for lipid estimation.

### Fly rearing for lipid estimation

To understand the fractional levels of lipid levels, assay was performed after 33 generations of selection for increased resistant cold shock. On the 12th-day post egg collection, vials were randomly divided into two treatments – (a) cold shock and (b) no shock treatment. For both cold shock and no shock treatment, flies were transferred into an empty glass vial, and the cotton plug was pushed deep into the bottom, leaving one-third of the vial space for flies. After that, flies were subjected to either of the treatments following the protocol described in the previous report [1,2,5]. For each treatment, gender, and population, four sets of vials with each containing a density of 50 males or 50 females have been subjected to cold shock or no shock treatment. After the treatment, flies were immediately transferred to a Plexiglas cage containing a fresh food plate. Twenty-four hours post cold shock treatment, a fresh food plate was given to the flies for one hour to lay the stored eggs, and followed by this, and another fresh food plate was provided for four hours. These plates with eggs were incubated for 18 hours to hatch eggs, and later from this plate, 30 first instar larvae were collected with a wet paintbrush and cultured into vials with 6 mL of food. For a given population and treatment, ten such vials were set up. These vials were carefully monitored for eclosion. Flies were collected within two hours of eclosion and were immediately flash-frozen using liquid nitrogen. Fifty males and 50 females per treatment and per population were frozen. These flies were stored at -80°C until assayed for lipid content.

### Lipid estimation

This assay was performed to assess the lipid levels in FSB and FCB flies, after 50 generations of selection for increased resistance to cold shock. Lipid estimation was performed following the protocol described in [21] with minor modifications. Ten sets were made for each gender and treatment or population, and each set had a group of five flies. Each set of flies was transferred to a clean 2 mL microcentrifuge tube. Ten such replicates for a given treatment, gender or populations were set up. These tubes of flies were incubated at 65°C for 48 h, bodyweight for each group of five flies was recorded using a fine microbalance. These flies were transferred back to a microcentrifuge tube. Following this, 1.5 mL of diethyl ether was added to each tube. These tubes were placed on a shaker for mild agitation for 24 h at 25°C. The lipids are extracted out into the ether. After 24 h, Diethyl ether was discarded, and flies were again dried at 70°C for 12 h. Bodyweight after lipid extraction was measured using the fine microbalance. Absolute lipid content was calculated as the difference between body weight before and after lipid extraction. Lipid content per fly was calculated by dividing this difference by the number of flies per sample. Fractional lipid content was measured by dividing the lipid content by the bodyweight of the given sample. Fractional lipid content was analyzed using a three-factor mixed model analysis of variance (ANOVA) with selection regime (FSB vs. FCB) and treatment (Cold shock vs. no shock) as fixed factors crossed with the block (1-5) as random factors. Multiple comparisons using Tukey’s HSD were performed. All the analyses were done at α = 0.05 level of significance using JMP PRO 16.

### Rearing of fly for gene expression assays

To determine the transcript levels of *Hsps* and *Fst* genes in males of the FSB and FCB. This assay was performed after 50 generations of selection for increase resistance cold shock. On the 9-10th day post-egg collection, males were collected as virgins and were held in groups of ten per vial (10 vials per treatment). Therefore, each treatment for a given population had 100 males. Two days later, *i. e*. on the 12th-day post-egg collection, males were randomly divided into two treatment groups – a) Cold Shock and b) No-Shock, and treatment was given as described in Singh et al. 2021. Soon after the treatment, flies were transferred to a Plexiglas cage. The transcript level of genes was measured using qRT-PCR of the flash-frozen flies during the recovery time post cold shock, *i*.*e*., 4 h or 12 h. Transcript levels of genes were analyzed using the Nonparametric Kruskal-Wallis test with Dunnett’s multiple comparisons test. All the analyses were done at α = 0.05 level of significance using GraphPad Prism 9.

### RNA extraction

Flies were homogenized in TRI reagent (Sigma-Aldrich) using a motorized pestle. RNA extraction was performed following the protocol and protocol described in Litwinoff et al. 2018. Extracted RNA was suspended in 30µL of DEPC treated water. Remnants of genomic DNA contamination were removed from the sample using RNase-Free DNase digestion and followed by RNA quality, and quantity was checked using NanoDrop 2000 Spectrophotometer (Thermo Scientific). Absorbance was measured at 260 nm and 280 nm. Absorbance 260/280 (A260/280) was calculated, and samples with A260/280 > 1.9 were further used. An equal amount of RNA of all samples was converted into cDNA using reverse transcriptase enzyme and random hexamers (Superscript III First-Strand Synthesis kit). The obtained cDNA was diluted ten-fold and used to measure the gene expression at mRNA levels.

### Quantitative Real-Time PCR (qRT-PCR)

Transcript levels of *Hsp22, Hsp23, Hsp40, Hsp68, Hsp70Aa* (*Hsp70*), *Hsp83*, and *Fst* genes were measured. Expression of a housekeeping gene *Actin* or *ribosomal protein S20* (*Rps20)* was used as an internal control. Primer sequences for all the genes were given in Table 2. Transcript levels of genes were measured using qRT-PCR with Fast SyBr green (Thermo Scientific) on Eppendorf Mastercycler. All the samples were run in duplicates. The cycle threshold (C_T_) values were obtained, and the expression of each gene of interest was normalized using the expression of internal control. The following calculations were done:

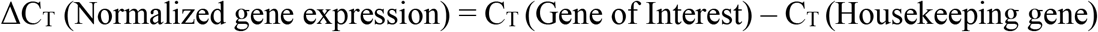

ΔC_T_ values under cold-shocked and non-shocked conditions were obtained for FSB and FCB. These ΔC_T_ values were further used to derive gene expression differences between FSB and FCB under either shocked or non-shocked conditions. Therefore, for all genes, cold-shocked/no shock FSB or cold-shock FCB was compared with no-shock FCB. This was denoted as ΔΔ C_T_ following:

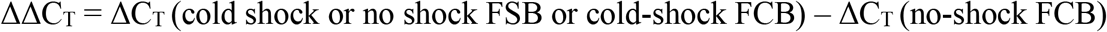

The difference in normalized gene expression between FSB and FCB (ΔΔ C_T_) was used to calculate the fold change in the expression of genes (Pfaffl 2001).

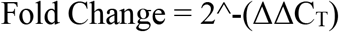

For the analysis, mean fold change across five blocks was calculated relative to the FCB no shock, expression for the given gene between FSB and FCB. The difference in normalized gene expression between FSB and FCB (ΔΔ C_T_) was used to calculate the fold change in the expression of genes follow previous report [22].

## RESULTS

### Transcript levels of genes

Transcript levels of *Hsp22, Hsp23, Hsp40, Hsp68, Hsp70Aa* (*Hsp70*), *Hsp83*, and *Fst* were assessed 4 h or 12 h post Cold shock (∼5°C) or No shock (25°C) (Fig. 1A-G). Expression of *Hsp22, Hsp23, Hsp40, Hsp68*, and *Fst* genes that we assayed was significantly higher under Cold-shock than No-shock conditions in both the FSB and FCB populations (data not shown), indicating that these genes are associated with cold shock response. However, the study aims to compare the expression of these genes between the FSB populations and FCB populations to address questions about the evolution of gene expression patterns at mRNA levels. The pattern was similar across the two-time points (4 h or 12 h) of recovery after cold shock. These results indicate no substantial difference in the expression of these genes between FSB and FCB populations.

**Figure 1:**
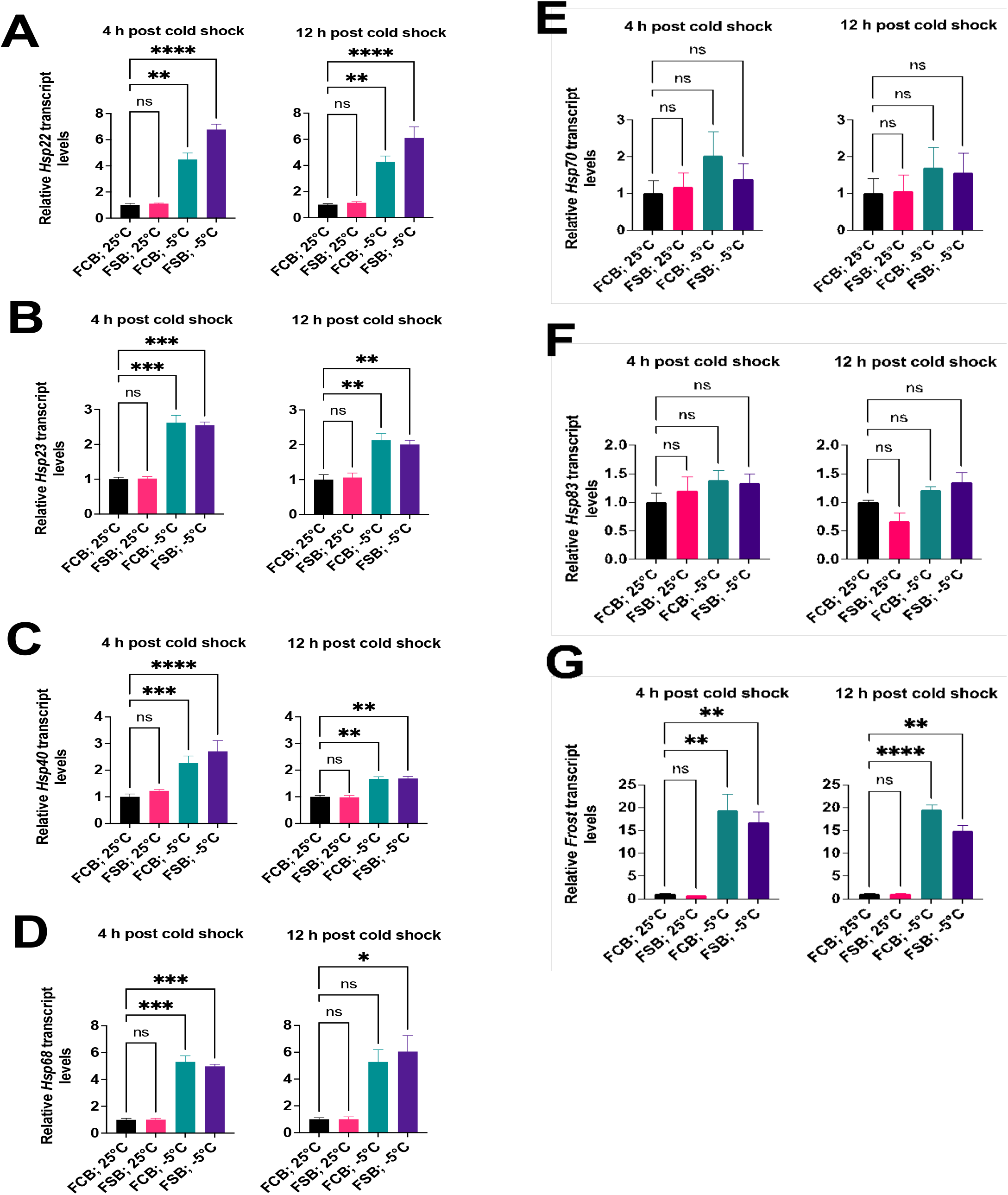
*Expression of Hsps and Fst* up regulated post cold shock treatment. **(A-G)** Relative transcripts levels of *Hsp22* **(A)**, *Hsp23* **(B)**, *Hsp40* **(C)**, *Hsp68* **(D)**, (*Hsp70* **(E)**, *Hsp83* **(F***), and* **(G)** *Fst* were measured at 4 h or 12 h post cold shock or no shock treatment to the FSB and FCB populations. Transcript levels of *Hsp22, Hsp23, Hsp40, Hsp68, and Fst were significantly in both the FSB and FCB populations at 4 h or 12 h post cold shock compared to no shock treatment*. qRT-PCR values were normalized to *Actin* and plotted relative to FCB no-shock. Statistical significance (Fig. **A-G**) was evaluated by Nonparametric, Kruskal-Wallis test with Dunnett’s multiple comparisons test, Data are represented as means ± SEM in bar graphs from different experiment. *p < 0.05, **p < 0.01, ***p < 0.

### Lipid content

We did not find a significant selection effect on fractional lipid content in both males (Table 1; Fig. 2A) and females (Table 1; Fig. 2B). A significant effect of treatment (Cold-shock vs. No-shock) was found in females (p = 0.038). Lower fractional lipid content under the No-shock condition was observed in females. None of the other interactions were found to be significant.

001.

**Table 1:**
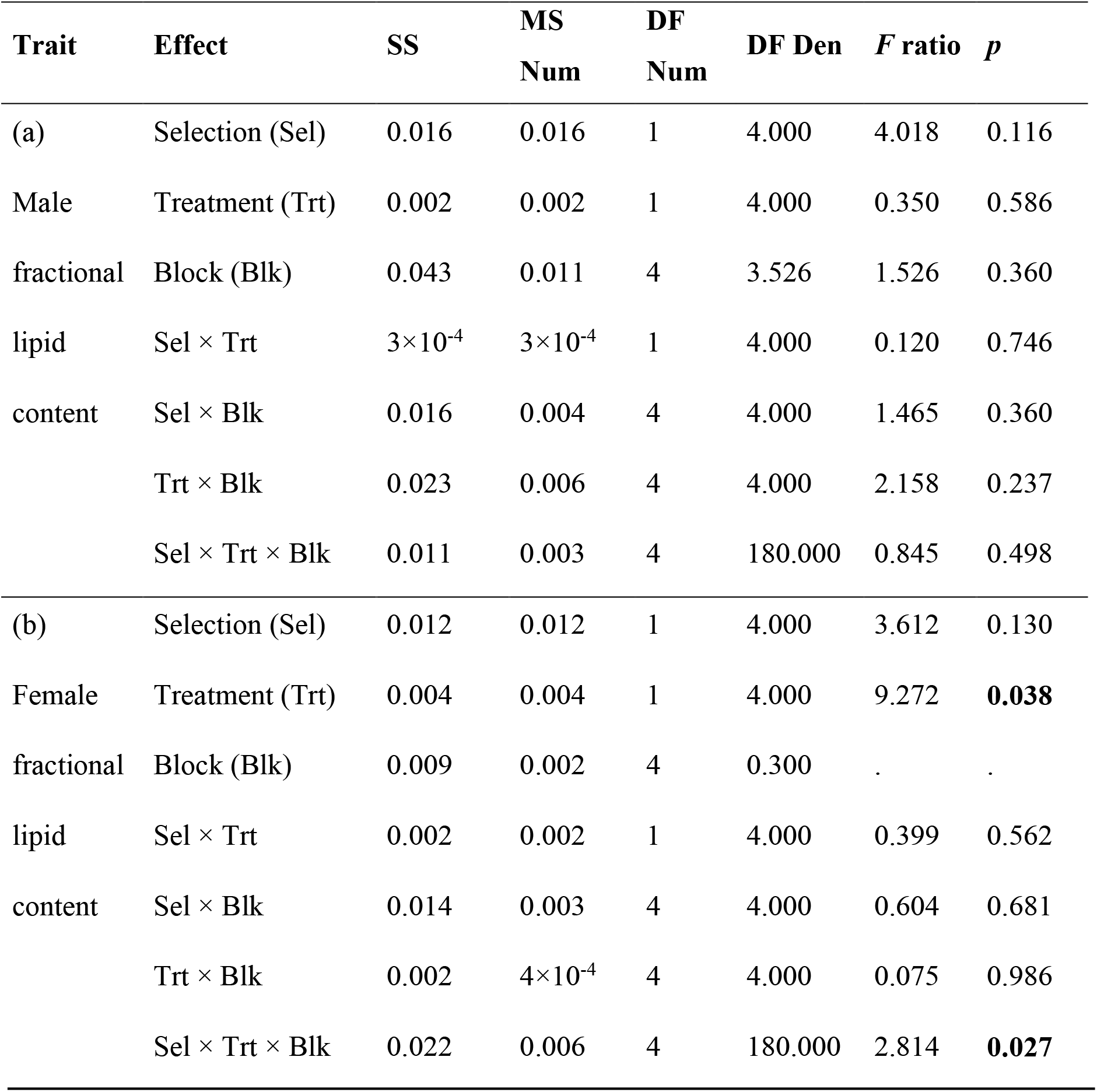
Summary of the results from the three-factor mixed model ANOVA on fractional lipid content in (a) male, and (b) female with selection regime (FSB and FCB) and treatment (cold shock and no shock) as fixed factors crossed with blocks (1-5) as a random factor. p-values in the bold case are statistically significant. Estimated denominator DF (Satterthwaite method) was very low. Hence F ratio and *p* values are unavailable for this effect.

**Table 2:**
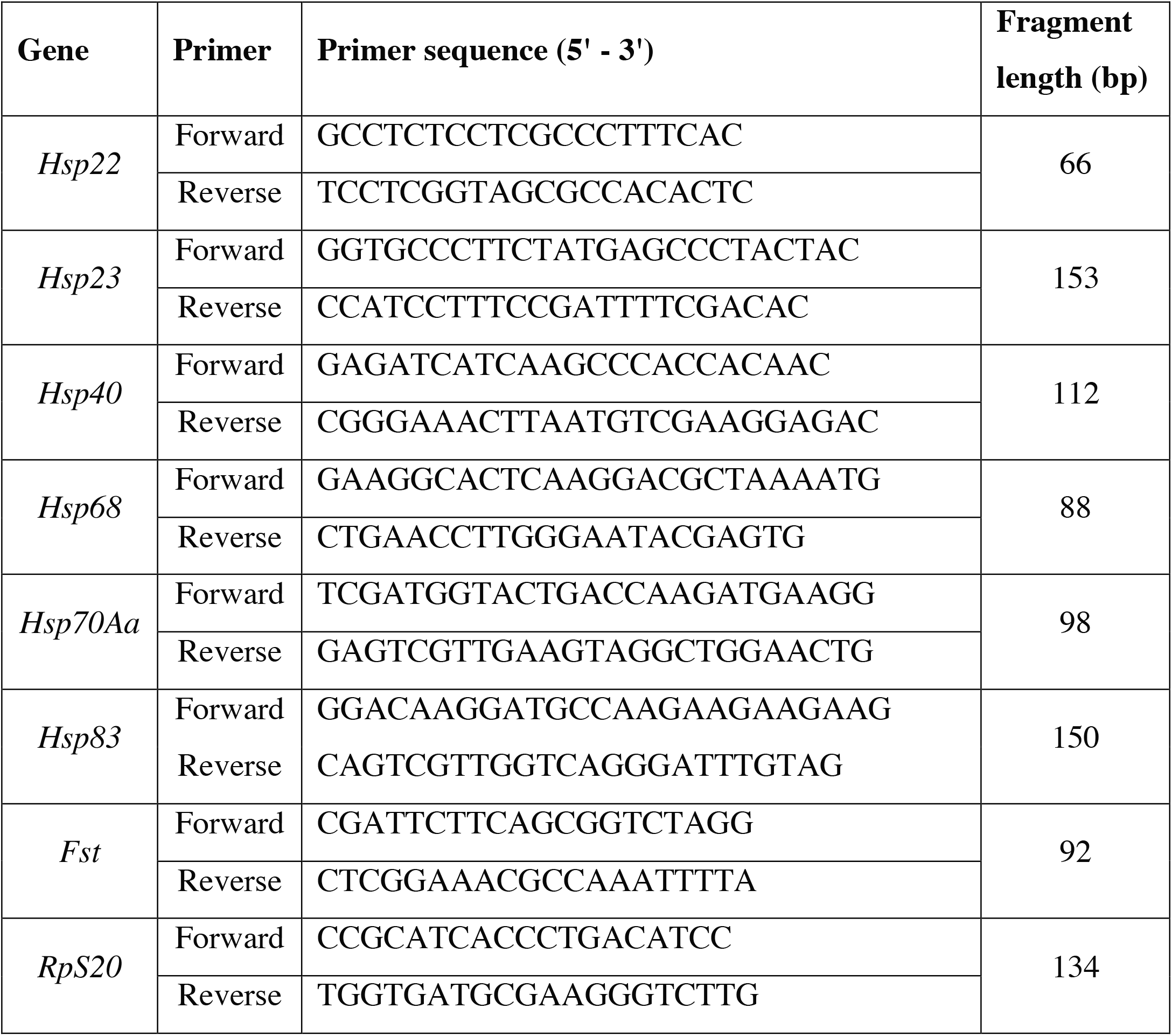
Primer sequences for genes used in this study.

**Figure 2:**
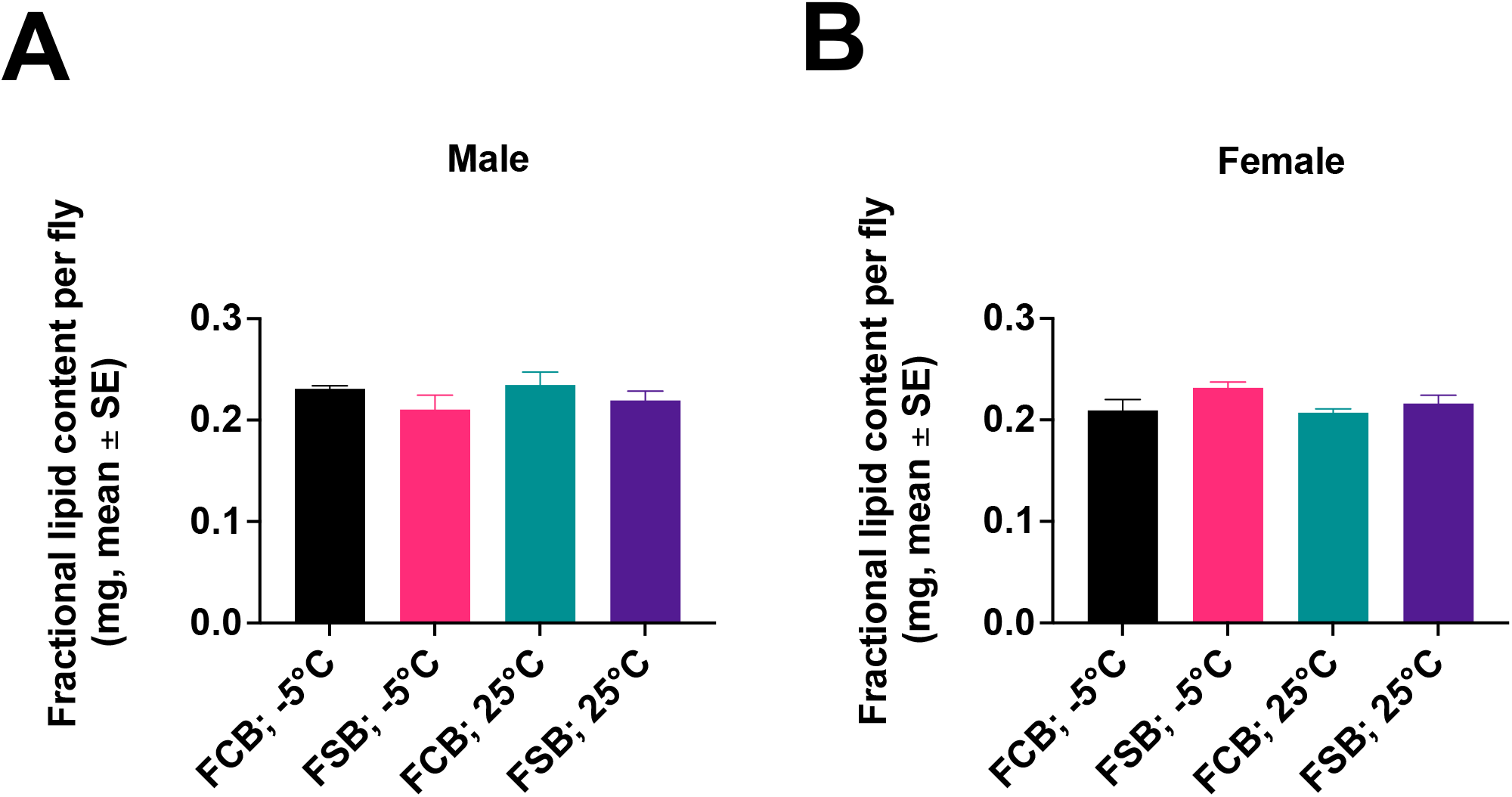
Effect of cold shock on the fractional lipid levels. **(A)** Fractional lipid content in the males of the FSB and FCB populations was assessed under the cold shock or no shock condition. We did not find significant effects of selection, treatment or selection × treatment interaction. Open bars represent FSB and closed bars represent FCB populations. (**B)** Similarly fractional lipid content was measured in the females of the both FSB and FCB population under the cold shock or no shock conditions. We did not find significant effects of selection or selection × treatment interaction. Open bars represent FSB, and closed bars represent FCB populations. Statistical significance (Fig. **A** and **B**) was evaluated using three-factor mixed model ANOVA on fractional lipid content in (**A**) male, and (**B**) female with selection regime (FSB and FCB) and treatment (cold shock and no shock) as fixed factors crossed with blocks (1-5) as a random factor. Data are represented as means ± SEM in bar graphs from different experiment. *p < 0.05, **p < 0.01, ***p < 0.001.

## DISCUSSION

The study investigated the underlying mechanisms of evolved cold tolerance. To measure these mechanisms, we examined (a) transcript level of several *Hsp22, Hsp23, Hsp40, Hsp68, Hsp70Aa* (*Hsp70*), *Hsp83*, and *Fst* genes under the cold shock or the benign condition, and (b) lipid content (b). This study presents the evidence of heightened transcripts levels of *Hsps* and *Fst genes* measured at 4 h or 12 h of recovery post cold shock compared to no shock conditions. However, no significant alteration in transcript levels of *Hsps* or *Fst gene* between FSB and FCB populations. This study reports the higher levels of lipid in the cold shock females of the both FSB and FCB population compared to no shock condition.

A number of physiological and biochemical changes are expected to occur when organisms experience stress, whether biotic or abiotic. Different kinds of stresses can elicit different responses. Cold stress often causes damage to proteins, and lipids in the cellular machinery. [23] showed that lipids play a significant role in maintaining cold tolerance in different species of *Drosophila*. This study did not find a difference in total lipid content between selected and control populations. However, we did not investigate the types of lipid species. Previous reports suggested that an increase in the unsaturated fatty acid content protects against cold stress [24]. Therefore, it’s possible that type of lipid might have evolved in the FSB populations without significantly changing fractional lipid content.

Heat shock proteins play a crucial role in mitigating the damage. We assessed the expression of several *Hsp22, Hsp23, Hsp40, Hsp68, Hsp70Aa* (*Hsp70*), *Hsp83*. In addition to *Hsps*, we also examined expression patterns of the *Fst* gene. *Fst* is linked with cold adaptation even though its molecular function is yet to be determined [25,26]. The expression of all the genes that were assayed was heightened under cold shocked conditions compared to non-shocked conditions suggesting that these genes are involved in the cold shock response. This finding agrees with the results of previous reports [25,26]. However, we did not find a significant change in transcript levels of either *Hsps* or *Fst* in the FSB populations relative to FCB populations. These results are contrary to the previous studies [26,27]. There could be various reasons for the observed results. Firstly, our selection maintenance protocol is different from [26]. Secondly, the time (post-recovery) at which measured the expression of these genes. Expression of *Hsps* is shown to be temporally regulated [25,26,27]). Colinet et al. [26] found differences in gene expression when measured after 2 h or 4 h of recovery. We selected the time points - 4 h, and 12 h during recovery post cold shock since the previous report documented the apparent differences in behavioral and fitness-related traits between the FSB and FCB populations at these time points. Therefore, there is a possibility that measurement of gene expression at different time points during and post-treatment might show differences in the gene expression. An additional point of caution is that we assayed only the levels of mRNA, not the functional proteins. It is quite possible that the levels of the proteins of these genes could be higher in the FSB populations even when the RNA levels are not different between FSB and FCB populations.

There are several ways organisms cope with stressful conditions. Further investigation into these mechanisms would better answer the mechanistic basis of evolved cold resistance in FSB populations. Future research into lipid composition (saturated vs. unsaturated lipids), other metabolites like glycogen, glycerol and proline [13,19] could help us find a clear answer. Microarray profiles [27,28,29]) of other cold-adapted lines in *D. melanogaster* have found a large number of other genes to be involved in cold stress. We also propose a future investigation into these genes to understand the mechanistic basis of the evolved response.

## Ethics declarations

### Ethics approval and consent to participate

Not applicable.

### Consent for publication

Not applicable.

### Availability of data and materials

All data analyzed or used for this study are available from the DRYAD database.

### Competing interests

All the authors declare no competing interests.

### Funding

This work was supported by the Indian Institute of Science Education and Research Mohali, Govt. of India.

## Contributions

KS and NGP conceived the experimental evolution regimes and associated protocols. KS: Conceptualized and designed the study, performed the experiments, analyzed the data, wrote and edited the manuscript. NGP: edited the manuscript, contributed reagents, materials, and analysis tools. All authors have read and approved the final version of the manuscript.

## Acknowledgements

Karan Singh thanks to Indian Institute of Science Education and Research Mohali, Govt. of India for the Junior and Senior Research Fellowship. We also thank Anand Rai for his assistance in the RT-PCR experiments.

## Author information

**Karan Singh**

Present address: Department of Cell Biology, NYU Grossman School of Medicine, 650 Medical Science Building, 550 First Ave, New York, NY, 10016, USA

## Affiliations

**Indian Institute of Science Education and Research Mohali, Knowledge City, Sector 81, SAS Nagar, PO Manauli, Mohali, Punjab, 140306, India**

Karan Singh & Nagaraj Guru Prasad

## Abbreviations

FSB: Freeze shock selected line derived from the BRB populations
FCB: Freeze shock control populations derived from the BRB populations
BRB: Blue Ridge Baseline populations
ANOVA: Analysis of variance
h: Hour

## Notes

### Competing Interest Statement

The authors have declared no competing interest.

### Summary of Updates

This was modifed in order to submit to BMC Ecology and Evolution

## References

1. Sinclair BJ, Ferguson LV, Salehipour-shirazi G, MacMillan HA. Cross-tolerance and cross-talk in the cold: relating low temperatures to desiccation and immune stress in insects. Integr Comp Biol. 2013;53: 545–556.

2. Singh K, Kochar E, Prasad NG. Egg Viability, Mating Frequency and Male Mating Ability Evolve in Populations of Drosophila melanogaster Selected for Resistance to Cold Shock. PLoS One. 2015;10: e0129992.

3. Singh K, Prasad NG. Evolution of pre- and post-copulatory traits in female Drosophila melanogaster as a correlated response to selection for resistance to cold stress. J Insect Physiol. 2016;91-92: 26–33.

4. Singh K, Samant MA, Tom MT, Prasad NG. Evolution of Pre- and Post-Copulatory Traits in Male Drosophila melanogaster as a Correlated Response to Selection for Resistance to Cold Stress. PLoS One. 2016;11: e0153629.

5. Singh K, Kochar E, Gahlot P, Bhatt K, Prasad NG. Evolution of reproductive traits have no apparent life-history associated cost in populations of Drosophila melanogaster selected for cold shock resistance. BMC Ecology and Evolution. 2021. doi:10.1186/s12862-021-01934-2

6. Chapman RF, Chapman RF. The Insects: Structure and Function. Cambridge University Press; 1998.

7. Bale JS. Insects and low temperatures: from molecular biology to distributions and abundance. Philosophical Transactions of the Royal Society of London. Series B: Biological Sciences. 2002. pp. 849–862. doi:10.1098/rstb.2002.1074

8. Gullan PJ, Cranston PS. The Insects: An Outline of Entomology. John Wiley & Sons; 2014.

9. Gulevsky AK, Relina LI. Molecular and genetic aspects of protein cold denaturation. Cryo Letters. 2013;34: 62–82.

10. Ronges D, Walsh JP, Sinclair BJ, Stillman JH. Changes in extreme cold tolerance, membrane composition and cardiac transcriptome during the first day of thermal acclimation in the porcelain crab Petrolisthes cinctipes. J Exp Biol. 2012;215: 1824–1836.

11. Feder ME, Hofmann GE. Heat-shock proteins, molecular chaperones, and the stress response: evolutionary and ecological physiology. Annu Rev Physiol. 1999;61: 243–282.

12. Nadeau D, Corneau S, Plante I, Morrow G, Tanguay RM. Evaluation for Hsp70 as a biomarker of effect of pollutants on the earthworm Lumbricus terrestris. Cell Stress Chaperones. 2001;6: 153– 163.

13. Cold-shock and chilling tolerance in Drosophila. J Insect Physiol. 1994;40: 661–669.

14. Hodkova M, Hodek I. Photoperiod, diapause and cold-hardiness. European Journal of Entomology. 2004. pp. 445–458. doi:10.14411/eje.2004.064

15. Teets NM, Peyton JT, Ragland GJ, Colinet H, Renault D, Hahn DA, et al. Combined transcriptomic and metabolomic approach uncovers molecular mechanisms of cold tolerance in a temperate flesh fly. Physiol Genomics. 2012;44: 764–777.

16. Dennis AB, Dunning LT, Sinclair BJ, Buckley TR. Parallel molecular routes to cold adaptation in eight genera of New Zealand stick insects. Sci Rep. 2015;5: 13965.

17. MacMillan HA, Sinclair BJ. The role of the gut in insect chilling injury: cold-induced disruption of osmoregulation in the fall field cricket, Gryllus pennsylvanicus. J Exp Biol. 2011;214: 726–734.

18. Shang Q, Pan Y, Peng T, Yang S, Lu X, Wang Z, et al. PROTEOMICS ANALYSIS OF OVEREXPRESSED PLASMA PROTEINS IN RESPONSE TO COLD ACCLIMATION IN Ostrinia furnacalis. Arch Insect Biochem Physiol. 2015;90: 195–208.

19. Koštál V, Korbelová J, Rozsypal J, Zahradníčková H, Cimlová J, Tomčala A, et al. Long-Term Cold Acclimation Extends Survival Time at 0°C and Modifies the Metabolomic Profiles of the Larvae of the Fruit Fly Drosophila melanogaster. PLoS ONE. 2011. p. e25025. doi:10.1371/journal.pone.0025025

20. Southon TE, Gehrken U. Effect of temperature on cold-hardiness and tissue ice formation in the adult chrysomelid beetle Melasoma collaris L. J Insect Physiol. 1997;43: 587–593.

21. Zwaan BJ, Bijlsma R, Hoekstra RF. On the developmental theory of ageing. I. starvation resistance and longevity in Drosophila melanogaster in relation to pre-adult breeding conditions. Heredity. 1991;66 (Pt 1): 29–39.

22. Litwinoff EMS, Gold MY, Singh K, Hu J, Li H, Cadwell K, et al. Myeloid ATG16L1 does not affect adipose tissue inflammation or body mass in mice fed high fat diet. Obes Res Clin Pract. 2018;12: 174–186.

23. Djawdan M, Chippindale AK, Rose MR, Bradley TJ. Metabolic Reserves and Evolved Stress Resistance in Drosophila melanogaster. Physiological Zoology. 1998. pp. 584–594. doi:10.1086/515963

24. Ohtsu T, Katagiri C, Kimura MT, Hori SH. Cold adaptations in Drosophila. Qualitative changes of triacylglycerols with relation to overwintering. J Biol Chem. 1993;268: 1830–1834.

25. Sinclair BJ, Gibbs AG, Roberts SP. Gene transcription during exposure to, and recovery from, cold and desiccation stress in Drosophila melanogaster. Insect Mol Biol. 2007;16: 435–443.

26. Colinet H, Lee SF, Hoffmann A. Temporal expression of heat shock genes during cold stress and recovery from chill coma in adult Drosophila melanogaster. FEBS J. 2010;277: 174–185.

27. Qin W, Neal SJ, Robertson RM, Westwood JT, Walker VK. Cold hardening and transcriptional change in Drosophila melanogaster. Insect Mol Biol. 2005;14: 607–613.

28. Telonis-Scott M, Hallas R, McKechnie SW, Wee CW, Hoffmann AA. Selection for cold resistance alters gene transcript levels in Drosophila melanogaster. J Insect Physiol. 2009;55: 549–555.

29. Zhang J, Marshall KE, Westwood JT, Clark MS, Sinclair BJ. Divergent transcriptomic responses to repeated and single cold exposures in Drosophila melanogaster. J Exp Biol. 2011;214: 4021–4029.

